# SARS-CoV-2 receptor binding domain fusion protein efficiently neutralizes virus infection

**DOI:** 10.1101/2021.04.18.440302

**Authors:** Abigael Chaouat, Hagit Achdout, Inbal Kol, Orit Berhani, Gil Roi, Einat B. Vitner, Sharon Melamed, Boaz Politi, Eran Zahavy, Ilija Brizic, Tihana Lenac Rovis, Or Alfi, Dana Wolf, Stipan Jonjic, Tomer Israely, Ofer Mandelboim

**Affiliations:** The Concern Foundation Laboratories at the Lautenberg Center for Immunology and Cancer Research, Institute for Medical Research Israel Canada (IMRIC), The Hebrew University Hadassah Medical School, Jerusalem, Israel; Israel Institute for Biological Research (IIBR), Ness-Ziona, Israel. Department of Infectious Diseases, Ness-Ziona 74100, POB 019, Israel; Department of Biochemistry and Molecular Genetics, Israel Institute for Biological Research, Ness Ziona, Israel; Center for Proteomics, Faculty of Medicine, University of Rijeka, Rijeka, Croatia; Lautenberg Center for General and Tumor Immunology, The Hebrew University Faculty of Medicine, Jerusalem, Israel; Clinical Virology Unit, Hadassah Hebrew University Medical Center, Jerusalem, Israel

## Abstract

Severe acute respiratory syndrome coronavirus 2 (SARS-CoV-2) is responsible for the COVID-19 pandemic, causing health and economic problems. Currently, as dangerous mutations emerge there is an increased demand for specific treatments for SARS-CoV-2 infected patients. The spike glycoprotein on the virus membrane binds to the angiotensin converting enzyme 2 (ACE2) receptor on host cells through its receptor binding domain (RBD) to mediate virus entry. Thus, blocking this interaction may inhibit viral entry and consequently stop infection. Here, we generated fusion proteins composed of the extracellular portions of ACE2 and RBD fused to the Fc portion of human IgG1 (ACE2-Ig and RBD-Ig, respectively). We demonstrate that ACE2-Ig is enzymatically active and that it can be recognized by the SARS-CoV-2 RBD, independently of its enzymatic activity. We further show that RBD-Ig efficiently inhibits in vitro and in vivo SARS-CoV-2 infection, better than ACE2-Ig. Mechanistically we show that anti-spike antibodies generation, ACE2 enzymatic activity and ACE2 surface expression were not affected by RBD-Ig. Finally, we show that RBD-Ig is more efficient than ACE2-Ig at neutralizing high virus concentration infection. We thus propose that RBD-Ig physically blocks virus infection by binding to ACE2 and that RBD-Ig should be used for the treatment of SARS-CoV-2-infected patients.

**Author Summary:** SARS-CoV-2 infection caused serious socio-economic and health problems around the globe. As dangerous mutations emerge, there is an increased demand for specific treatments for SARS-CoV-2 infected patients. SARS-CoV-2 infection starts via binding of SARS-CoV-2 spike protein receptor binding domain (RBD) to its receptor, ACE2, on host cells. To intercept this binding, we generated Ig-fusion proteins. ACE2-Ig was generated to possibly block RBD by binding to it and RBD-Ig to block ACE2. We indeed showed that the fusion proteins bind to their respective target. We found that it is more efficient to inhibit SARS-CoV-2 infection by blocking ACE2 receptor with RBD-Ig. We also showed that RBD-Ig does not interfere with ACE2 activity or surface expression. Importantly, as our treatment does not target the virus directly, it may be efficient against any emerging variant. We propose here that RBD-Ig physically blocks virus infection by binding to ACE2 and thus it may be used for the treatment of SARS-CoV-2-infected patients.

## Main text

### Introduction

SARS-CoV-2 was first reported in December 2019 in China. It is a highly contagious virus which had caused worldwide socio-economic, political, and environmental problems [1]. In an attempt to stop the pandemic, the FDA first issued an emergency use authorization for Pfizer [2] and Moderna [3] vaccines, followed by Ad26.COV2.S [4]. Both vaccines, The Pfizer vaccine called BNT162b2 [5], and Moderna vaccine called mRNA-1273 [6], are composed of a lipid-nanoparticle (LNP)–encapsulated mRNA expressing the prefusion-stabilized spike glycoprotein. However, a treatment that will inhibit virus infection is urgently needed because not all individuals will be vaccinated, and even in those that will, the vaccines are not 100% effective. Furthermore, lately dangerous virus mutants appeared which may affect vaccine efficiency [7].

To infect cells, the spike glycoprotein, located on SARS-CoV-2 envelope, binds to the ACE2 receptor found on host cells [8]. The spike protein is trimeric, where each monomer contains two subunits: S1 and S2, which mediate attachment and membrane fusion, respectively. S1 itself can be subdivided further into S1a and S1b, where the latter includes the RBD [9]. The virus binds primarily to ACE2 receptors on type 2 pneumocytes [10], thus it mainly targets the lungs, but as ACE2 is present on many other cells, it is also capable of causing damage to other organs such as the heart, the liver, the kidneys, blood and immune system [11]. ACE2 is a carboxypeptidase of the renin-angiotensin hormone system that is a critical regulator of blood volume, systemic vascular resistance, and thus cardiovascular homeostasis [12]. ACE2 converts angiotensin I to angiotensin 1-9, a peptide with anti-hypertrophic effects in cardiomyocytes [13], and angiotensin II to angiotensin 1-7, which acts as a vasodilator [14].

SARS-CoV-2 life cycle starts with its RBD binding to the ACE2 receptor and ends by release of virions which binds to ACE2 receptors elsewhere [10]. Thus, intercepting the binding of the virions to the ACE2 receptor may help to treat infection. Developing treatment for SARS-CoV-2 infection is especially important since the FDA has yet approved any specific treatment for SARS-CoV-2 infected patients [15].

To intercept SARS-CoV-2 RBD binding to ACE2 we have generated fusion proteins containing the extracellular portions of RBD and ACE2 which are fused to the Fc portion of human IgG1. We have chosen this approach since the Fc partner increases the half-life of the protein and enables efficient purification [16]. Indeed, using the IgG Fc as a fusion partner to significantly increase the half-life of a therapeutic peptide or protein was first described in 1989 [17]. Since then, Fc‐ fusion proteins have been investigated for their effectiveness to treat many pathologies. Most Fc‐ fusions target receptor- ligand interactions and thus are used as antagonists to block receptor binding (e.g. Etanercept, Aflibercept, Rilonacept, Belatacept, Abatacept) [18]. It has been shown that soluble extracellular domains of ACE2 can act as a decoy, competitive inhibitors for SARS-CoV-2 infection [19,20]. RBD-Ig, on the other hand was tested only as a preventive vaccine against SARS-CoV-2 and not as a possible treatment during active infection [21,22]. Taken together, we decided to assess whether fusion proteins consisting of either ACE2 or RBD could potentially serve as therapeutics for treating active SARS-CoV-2 infection. Importantly, we demonstrate both in vitro and in vivo that RBD-Ig is more efficient than ACE2-Ig in its ability to inhibit SARS-CoV-2 infection. We demonstrated that RBD-Ig binding to ACE2 does not interfere with its expression on the cell surface or with its enzymatic activity and suggest that RBD-Ig inhibits SARS-CoV-2 infection by physically interacting with ACE2.

## Results

### Generation of ACE2-Ig and RBD-Ig

Since binding of SARS-CoV-2 RBD to ACE2 on host cells mediates virus infection ([8] and Figure 1A, left), we decided to intercept this binding. For that, we generated fusion proteins composed of the extracellular portions of human ACE2 or the viral RBD fused to the Fc portion of human IgG1. These fusion proteins are expected to inhibit SARS-CoV-2 infection by either blocking SARS-CoV-2 spike protein with ACE2-Ig (Figure 1A, middle) or by blocking ACE2 on host cells with RBD-Ig (Figure 1A, left). To investigate whether the RBD-Ig we have generated can indeed bind to ACE2, we expressed ACE2 with an N-terminal flag-tag in 293T cells (293T-ACE2). Expression of ACE2 was verified by flow cytometry using an anti-flag antibody (Figure 1B). We then stained 293T-ACE2 cells with RBD-Ig and demonstrated that RBD-Ig binds to these cells, but not to the parental 293T cells (Figure 1C).

**Figure 1:**
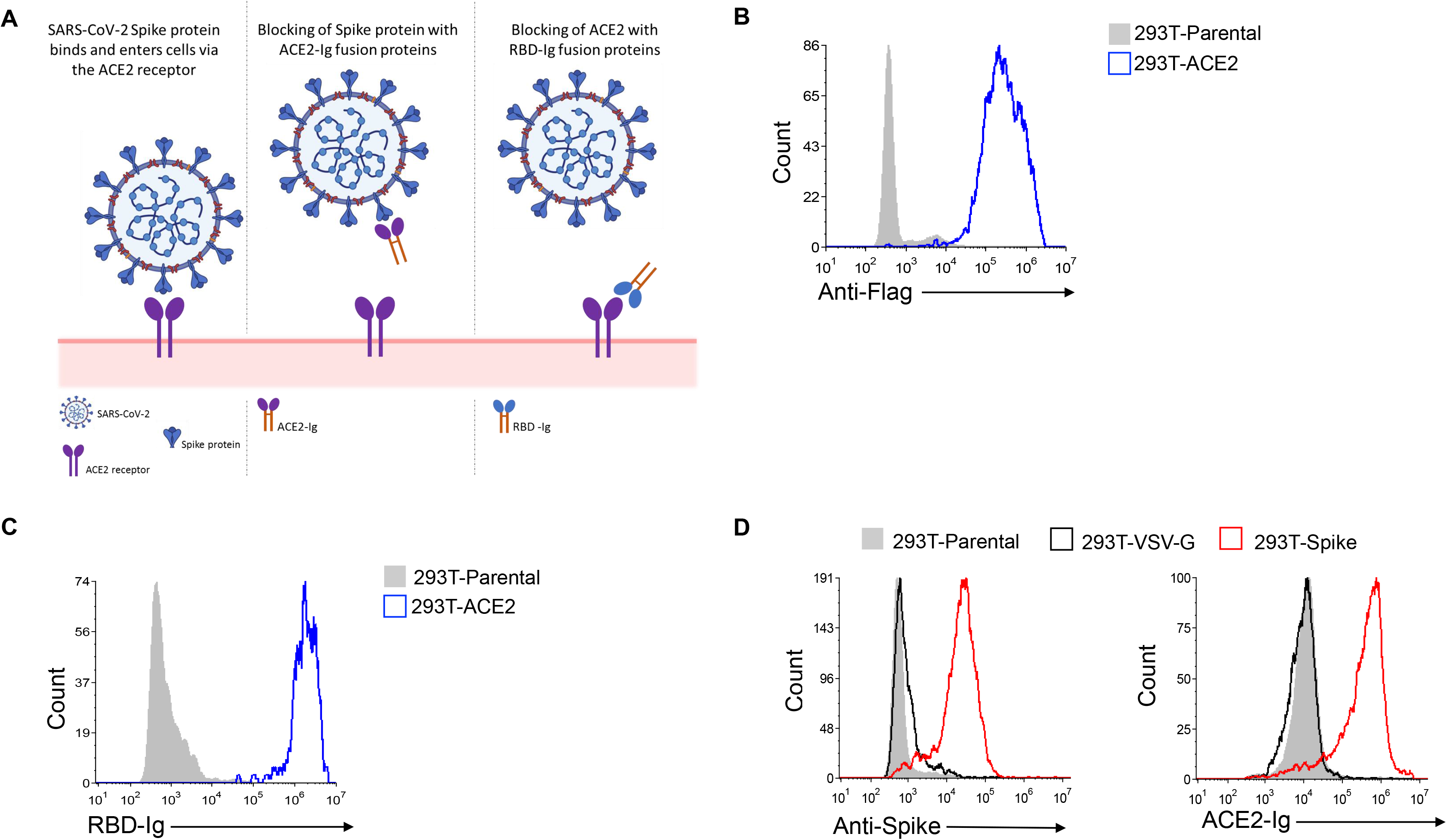
RBD-Ig and ACE2-Ig bind their respective target. (A) Schematic representation of our proposed treatments. SARS-CoV-2 infects ACE2 expressing cells (left panel). Binding of ACE2-Ig to SARS-CoV-2 Spike protein (middle panel) or binding of RBD-Ig to the ACE2 receptor (right panel) may prevent infection. (B) Staining of cells transfected to express ACE2 with an N-terminal Flag-tag (293T-ACE2 cells) and their parental cells that do not express a tag. This staining was performed using an anti-Flag antibody. (C) Staining of 293T-ACE2 cells with RBD-Ig. (D) Left panel: Spike protein surface expression on 293T cells co-transfected with either SARS-CoV-2 Spike envelope plasmid (293T-Spike cells) or Vesicular stomatitis virus (VSV) G envelope plasmid (293T-VSV-G cells). Right panel: Staining of 293T-Spike cells with ACE2-Ig. All histograms except from those made for 293T-Parental cells, were made from GFP positive gated cells. Figures shows one representative experiment out of 3 performed.

Next, to investigate the ability of ACE2-Ig to bind SARS-CoV-2 spike protein, we co-transfected 293T cells with SARS-CoV-2 spike envelope plasmid, a packaging plasmid and a GFP plasmid (293T-Spike). As a control we co-transfected 293T cells with a VSV-G envelope plasmid, a packaging plasmid and a GFP plasmid (293T-VSV-G). Staining was performed on GFP-positive gated cells. As can be seen, the 293T-Spike cells express high levels of the spike protein (Figure 1D, left), and were specifically recognized by ACE2-Ig. As expected, ACE2-Ig did not bind to the 293T-VSV-G cells (Figure 1D, right).

### ACE2 enzymatic activity is not required for its binding to SARS-CoV-2 spike protein

After confirming the binding of the fusion proteins to their respective targets, we wanted to check if ACE2-Ig is enzymatically active since enzymatic activity of ACE2 might be important for the course of COVID-19 disease [23,24]. To test the enzymatic activity, we used a commercial kit detailed in the “Methods” section. As can be seen in Figure 2A, ACE2-Ig was as active as human recombinant ACE2. Furthermore, the enzymatic activity of both proteins was completely abolished in presence of an ACE2 inhibitor (Figure 2A). After assessing ACE2-Ig activity, we wanted to test whether the enzymatic activity of ACE2 is required for its recognition by the SARS-CoV-2 spike protein. For that, we stained 293T-spike cells with ACE2-Ig in the presence or absence of an ACE2 inhibitor. We used a concentration of ACE2-Ig at which complete inhibition of enzymatic activity was achieved in the presence of the inhibitor (Figure 2A). As can be seen, no difference in ACE2-Ig binding was observed, regardless of whether the inhibitor was present or not (Figure 2B). Thus, we concluded that ACE2-Ig binding to SARS-CoV-2 spike protein is not dependent on its enzymatic activity.

**Figure 2:**
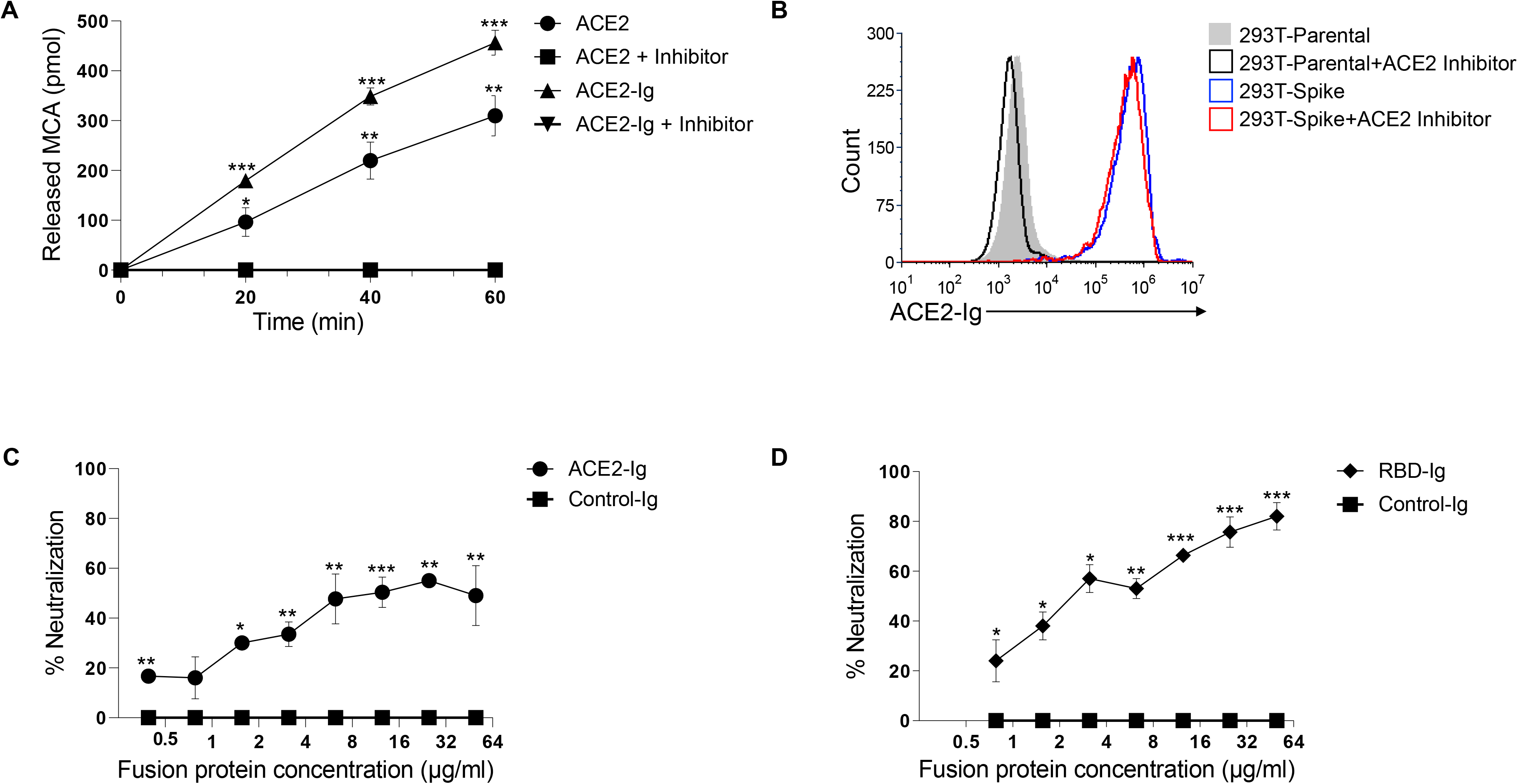
ACE2-Ig and RBD-Ig inhibits in vitro SARS-CoV-2 infection. (A) ACE2 enzymatic activity assay. Recombinant human ACE2 and ACE2-Ig were incubated with and without an ACE2 inhibitor, then MCA based peptide substrate was added and plate was immediately inserted in the fluorescent plate reader. *p<0.005, **p<0.0005, ***p<0.00005, Student’s t-test as compared to same treatment with inhibitor. (B) Staining of 293T-Spike cells with ACE2-Ig which was previously incubated for 15 minutes with or without an ACE2 inhibitor. (C-D) Plaque reduction neutralization test. Vero E6 cells were infected with SARS-CoV-2 and treated with increasing concentrations of either Control-Ig, ACE2-Ig (C) or RBD-Ig (D). % Neutralization was calculated as the percent of the decrease in plaque numbers, as compared with the background control. *P < 0.05; **P< 0.01; ***P <0.001; Student’s t-test as compared to Control-Ig. Figures shows one representative experiment out of 3 performed.

### RBD-Ig and ACE2-Ig inhibit SARS-CoV-2 infection in vitro

Next, we wanted to test whether in vitro SARS-CoV-2 infection can be inhibited by ACE2-Ig or RBD-Ig. To this end, we performed a plaque reduction neutralization test (PRNT), using Vero E6 cells that are permissive to SARS-CoV-2 infection [25]. Our negative control throughout these assays was a control fusion protein (Control-Ig). To assess inhibition by ACE2-Ig we initially incubated increasing concentration of the fusion protein with 300 PFU/ml of SARS-CoV-2 for 1 hour at 37°C, and then infected Vero E6 cells. Conversely, to test for inhibition by RBD-Ig, we had to first incubate the fusion protein with Vero E6 cells for 1 hour at 37°C, and then infect with 300 PFU/ml of SARS-CoV-2. Both strategies required a 48-hour incubation period to allow for plaque formation, followed by counting of said plaques and calculation of neutralization percentage. While the Control-Ig had no neutralizing effect in any of the concentrations tested, a dose-dependent neutralization of virus infection was observed for ACE2-Ig (Figure 2C), as well as for RBD-Ig (Figure 2D). When comparing between RBD-Ig and ACE2-Ig neutralization efficiency, RBD-Ig was significantly more efficient at the highest concentration used. When 50 ug/ml of RBD-Ig was applied, ~75% neutralization was observed (p=0.001) as compared to ~60% neutralization by ACE2-Ig. These results suggest that RBD-Ig inhibits in vitro infection to a greater degree.

### RBD-Ig efficiently inhibits SARS-CoV-2 infection in vivo

As RBD-Ig was more efficient than ACE2-Ig in vitro we wanted to further examine the fusion proteins efficiency in vivo. For that purpose, we infected homozygous female K18-hACE2 transgenic mice [26] by inhalation of 200 PFU of SARS-CoV-2. As a control for the infection, we looked at naïve (uninfected and untreated) mice. We also had a control for the treatment which included infected mice treated with an unrelated fusion protein (Control-Ig). The experiments lasted 15 days, during 1-5 days post-infection (dpi) the mice were injected three times intraperitoneally with 75ug of either RBD-Ig, ACE2-Ig, or Control-Ig. Treatments started 24 hours following infection.

Mice from the Control-Ig treated group started dying or losing more than 30% of their initial body weight (which is considered non ethical) at 7 dpi and we therefore could no longer use weight loss to assess the efficacy of our treatment. Thus, percentage of initial body weight was calculated until 7 dpi (Figure 3A). All SARS-CoV-2 infected mice started to lose weight at around 4 dpi. At 6-7 dpi the mice treated with RBD-Ig showed significantly less weight loss, as compared to all other infected groups (Figure 3A). We also monitored mice survival. Importantly, while the infected mice groups treated with Control-Ig or ACE2-Ig showed ~20% survival, the RBD-Ig treated group had significantly higher percentage with 50% survival (Figure 3B).

**Figure 3:**
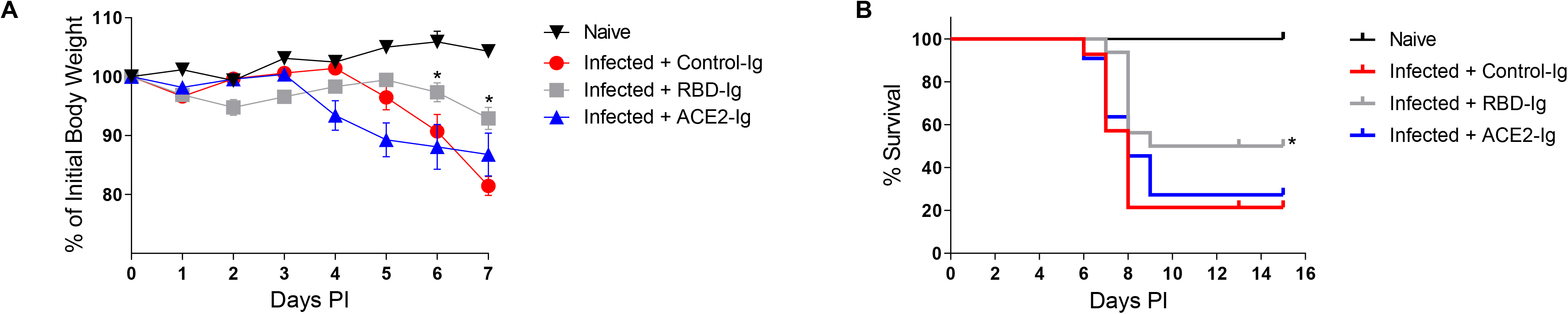
RBD-Ig decreases disease severity of SARS-CoV-2 infected mice. (A) Homozygous female K18-hACE2 transgenic mice were infected with SARS-CoV-2 (day 0) and treated with 75ug/mouse of either Control-Ig, RBD-Ig or ACE2-Ig. % of initial body weight was calculated from mice which were weighed daily. (B) Survival percentages of SARS-CoV-2 infected mice treated as described in A. *P < 0.05; Mantel-Cox test as compared to Infected + Control-Ig. Figure shows the combined results of two independent experiments.

### All infected mice generate neutralizing anti-Spike antibodies

Finally, we wanted to investigate why the ACE2-Ig fusion protein was less effective than RBD-Ig, both in vitro (Figure 2) and in vivo (Figure 3). We therefore evaluated anti-spike and anti-ACE2 IgG antibody generation in all mice groups. For that, sera were collected at 15 dpi from all mice groups including naïve mice and various sera dilutions were used to stain 293T-Spike cells and 293T-ACE2 cells to assess antibody existence and quantity by flow cytometry (Figure 4A). To our surprise, an equivalent, dose-dependent staining of all 293T-Spike cells was observed with sera obtained from all infected and treated mice (Figure 4A). As expected, no staining was observed when sera of naïve uninfected mice were used (Figure 4A) or when the sera from all mice groups were used to stain the 293T-ACE2 cells (Figure 4A). Thus, we concluded that the quantity of antibodies generated is not the reason for why RBD-Ig is more efficient. We then wanted to check the quality of the antibodies, as we suspected that RBD-Ig treated mice will generate more neutralizing antibodies since it was shown that anti-RBD antibodies have neutralization effect [27]. To test this hypothesis, we used the 293T-Spike cells and stained them with ACE2-Ig in the presence or absence of sera obtained from all mice groups. Since naïve mice did not generate anti-spike antibodies (Figure 4A), their sera, as expected, did not contained neutralizing antibodies. Indeed, similar ACE2-Ig binding was observed with and without blocking (Figure 4B). In all the infected mice, a comparable level of blocking was seen with sera regardless of the treatment administered, as assessed by reduced ACE2-Ig staining (Figure 4B). From these results we concluded that all the antibodies that were generated were a result of SARS-CoV-2 infection rather than our treatment.

**Figure 4:**
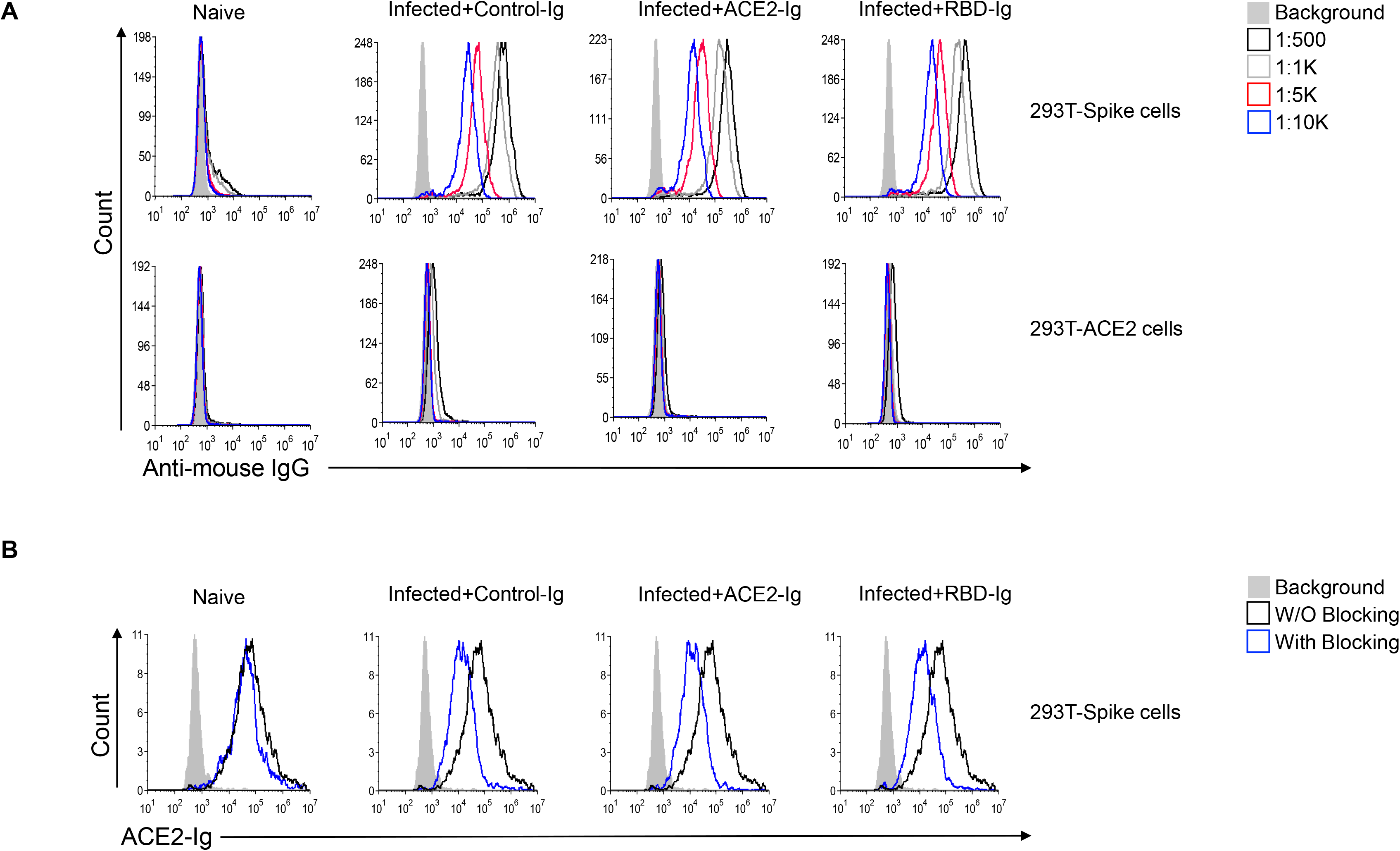
Blocking anti-Spike antibodies are generated in all immunized mice. (A) Anti-Spike IgG antibodies generated by mice following infection with SARS-CoV-2. Sera were taken 15 dpi from all mice groups and from naïve mice and diluted as indicated (upper right). Sera was incubated either with 293T-Spike cells (upper histograms) or with 293T-ACE2 cells (lower histograms) as a primary antibody then cells were stained with Alexa fluor 647 anti-mouse IgG secondary antibody. (B) ACE2-Ig staining of 293T-Spike cells in the presence or absence of sera from the various groups. Sera from all indicated groups were incubated with 293T-Spike cells for 1 hour at 4°C followed by staining with ACE2-Ig. All histograms were gated on GFP positive cells. Figure shows one representative experiment out of 2 performed.

### SARS-CoV-2 infection is possibly inhibited via physical blockade of ACE2 by RBD-Ig

To further investigate why is RBD-Ig better than ACE2-Ig at inhibiting SARS-CoV-2 infection we first generated a specific monoclonal antibody against ACE2 using ACE2-Ig as an antigen. This step was essential since the commercial antibodies (#ABIN1169449 and #MA5-32307) we tested did not recognize ACE2 effectively. As can be seen in Figure 5A, our generated antibody (anti-ACE2 01) is specific to ACE2, as it binds only to the 293T-ACE2 cells. To test if the antibody blocks the interaction with SARS-CoV-2 RBD, we incubated the antibody with 293T-ACE2 cells for 1 hour and then stained the cells with RBD-Ig. The anti-ACE2 01 antibody has no blocking property as its presence did not interfere with the binding of RBD-Ig to the ACE2 (Figure 5B).

**Figure 5:**
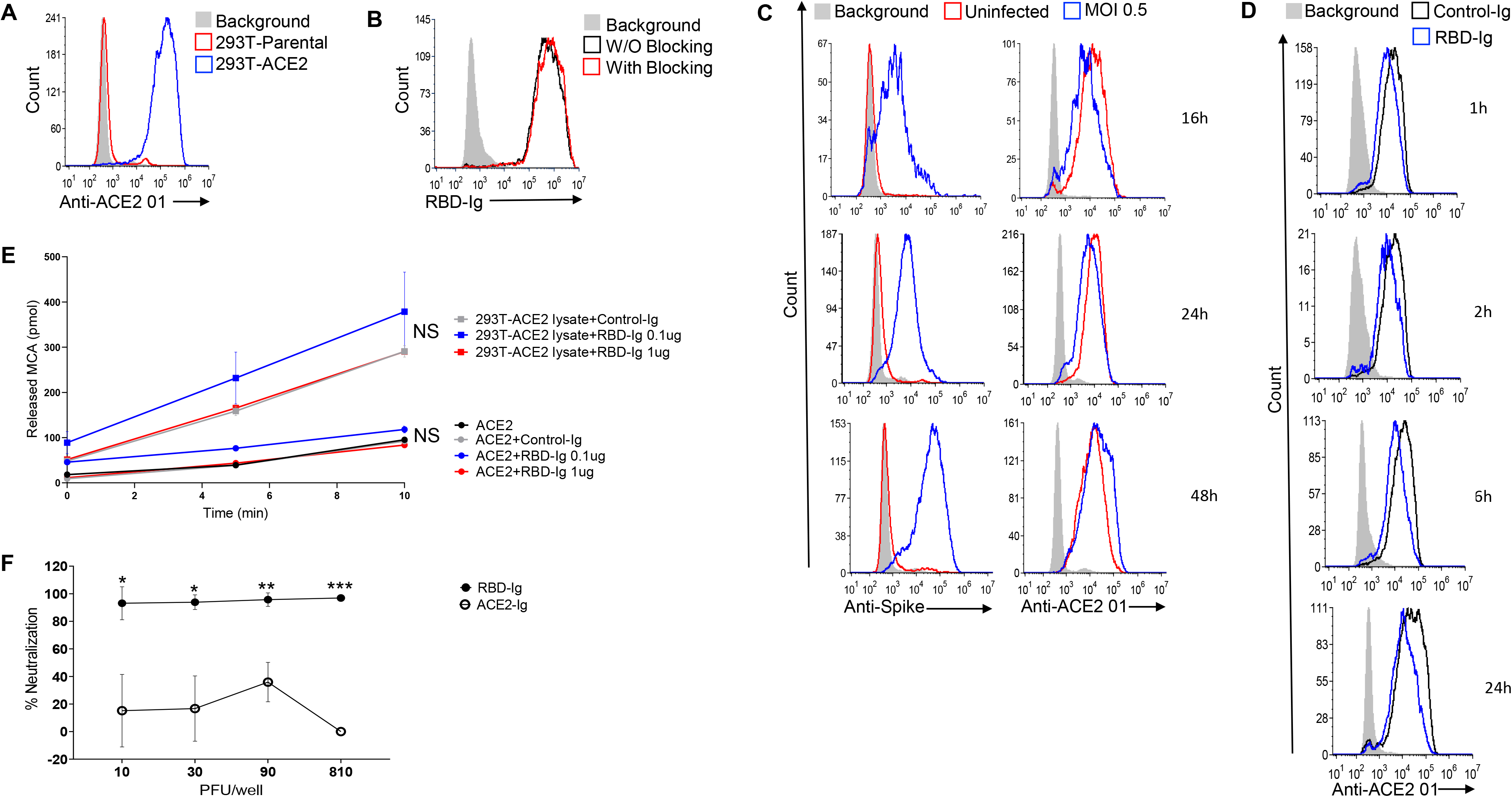
Effects of RBD-Ig. (A) Staining of 293T-Parental cells and 293T-ACE2 cells with the mAb anti-ACE2 01 we generated. (B) Staining of 293T-Parental cells and 293T-ACE2 cells with RBD-Ig. Cells were incubated with or without anti-ACE2 01 for 1 hour at 4°C, washed and then staining was performed. (C) Staining of infected (MOI 0.5) and uninfected VERO E6 cells with either an anti-Spike antibody to verify infection (left panel) or with our anti-ACE2 01 antibody (right panel) at 16,24,48 hours PI. (D) Staining with anti-ACE 01 of 293T-ACE2 cells which were incubated with 1 ug of either Control-Ig or RBD-Ig for 1,2,6 and 24 hours. (A-D) All histograms were gated on GFP positive cells. (E) ACE2 enzymatic activity assay. Recombinant human ACE2 and 293T-ACE2 cells lysate (10 ug) were incubated with either Control-Ig (1 ug) or RBD-Ig (0.1 ug or 1ug), then MCA based peptide substrate was added and plate was immediately read in the fluorescent plate reader. Not significant (NS), Student’s t-test as compared with Control-Ig. (F) Plaque reduction neutralization test. Vero E6 cells were infected with increasing SARS-CoV-2 titers and treated with 20 ug/well of either Control-Ig, ACE2-Ig or RBD-Ig. % Neutralization was calculated as the percent of the decrease in plaque numbers, as compared with cells treated with Control-Ig. *P < 0.01; **P< 0.005; ***P <0.00001; Student’s t-test. Figures shows one representative experiment out of 3 (A-E) or 2 (F) performed.

We then analyzed whether the expression of ACE2 is altered, at various time points, following SARS-CoV-2 infection. We infected 293T-ACE2 cells with a 0.5 MOI and compared between ACE2 surface expression on infected cells to uninfected cells using our anti-ACE2 01 antibody. Cells were harvested at 16-, 24- and 48-hour post-infection and SARS-CoV-2 spike surface expression was assessed by flow cytometry to verify infection. Infected cells indeed expressed SARS-CoV-2 spike protein while uninfected cells did not (Figure 5C, left). Little or no change in ACE2 surface expression was noticed at all the time points (Figure 5C, right), indicating that ACE2 surface levels are not subjected to changes following infection. We then tested if RBD-Ig incubation with 293T-ACE2 cells will lead to reduced ACE2 surface expression as we hypothesized that this might be the reason why RBD-Ig is more efficient than ACE2-Ig at neutralizing infection. We incubated RBD-Ig or Control-Ig with 293T-ACE2 cells for 1, 2, 6 and 24 hours. Following incubation cells were harvested and ACE2 surface expression was assessed by flow cytometry using our generated antibody, anti-ACE2 01. As can be seen in Figure 5D, ACE2 surface levels were only slightly reduced following RBD-Ig binding, suggesting that this is not the reason why RBD-Ig is superior to ACE2-Ig.

Next, we examined whether ACE2 activity will be altered following interaction with RBD-Ig, as we thought that maybe the activity of ACE2 might affect somehow the infection. We incubated 0.1 or 1 ug of RBD-Ig or Control-Ig with recombinant human ACE2 or with 293T-ACE2 cells lysate. ACE2 activity was not affected by RBD-Ig when incubated with human ACE2 or with a lysate containing ACE2 (Figure 5E). These combined results suggest that treatment with RBD-Ig inhibits infection without affecting ACE2 activity and surface levels expression.

Our last assumption was that RBD-Ig inhibits infection by physically blocking ACE2. We further hypothesized that RBD-Ig is more efficient than ACE2-Ig because RBD-Ig binds to ACE2, for which surface expression does not change following infection (Figure 5C, right). In contrast, ACE2-Ig may be less efficient since it targets the spike protein of a constantly replicating virus. To test this hypothesis, we performed a plaque reduction neutralization test (PRNT) as described above (Figure 2C and D). However, instead of increasing the fusion protein concentration, we used one concentration of the fusion proteins (20ug/well) and increased SARS-CoV-2 titers. Neutralization percentages were calculated as compared to respective Control-Ig. RBD-Ig treatment was significantly more efficient at inhibiting in vitro SARS-CoV-2 infection as compared to ACE2-Ig (Figure 5F).

## Discussion

Saturday 30 January 2021 marked one year since the WHO declared COVID-19 as an international concerning health emergency. At that time, only 9826 SARS-CoV-2 cases were reported in 20 countries. As of February 2, 2021, the total number of cases is ~102 million with ~2.2 million deaths reported in 222 countries [28]. As it is now clear that the pandemic will not end soon, a treatment for SARS-CoV-2 infected patients is urgently needed. The need arises since vaccination with the Moderna or Pfizer vaccines had just started, and even vaccinated individuals might not be fully protected. First as the vaccines efficiency is ~95% [29] and second because lately alarming SARS-CoV-2 spike mutations were observed in different countries and were transmitted across the world [30][31][32][33]. While the E484K mutations only reduces neutralizing activity of human convalescent and post-vaccination sera [34], the SARS-CoV-2 501Y.V2 south African variant contains multiple mutations that may enable escape from neutralizing antibodies [35]. These data suggest that reinfection with antigenically distinct variants is possible and may reduce efficacy of current spike-based vaccines.

Recently, Bamlanivimab, a recombinant, neutralizing human IgG1 monoclonal antibody against SARS-CoV-2 spike protein has been authorized by the FDA under an emergency use authorization [36]. But as antibodies are highly specific, there is a risk that the virus will develop escape mutations. This scenario is less likely when using a full protein or one of its domains. For that purpose, we generated the fusion proteins ACE2-Ig and RBD-Ig and tested their functionality. We also tested the ACE2-Ig enzymatic activity since it is known that dysregulation of ACE2 activity can adversely exacerbate lung inflammation and injury [37,38], and induce a general pro-inflammatory response [39,40]. After demonstrating that ACE2-Ig in enzymatically active, we wanted to examine whether ACE2 activity is required for the binding to SARS-CoV-2 spike protein. We demonstrated that the enzymatic activity of ACE2 is not required for its recognition by SARS-CoV-2 RBD. Confirming these results, it was previously reported that binding of SARS-CoV spike protein to ACE2 is also independent of ACE2 catalytic activity [41].

We showed that ACE2-Ig inhibits in vitro SARS-CoV-2 infection as it has been previously shown [42] and that RBD-Ig inhibits infection significantly more than ACE2-Ig. Furthermore, we show that treatment with RBD-Ig using SARS-CoV-2 K18-hACE2 infected mice led to decrease in disease severity as assessed by reduced body weight and increased mice survival. Importantly, 50% of the RBD-Ig treated mice survived although active infection occurred, while ACE2-Ig injection had no effect. We think that the reason behind the low efficiency of ACE2-Ig in vivo is due to the low concentrations of fusion protein we administered which was 75ug/mouse, injected intraperitoneally. Indeed, when Iwanaga et al injected intravenously ACE2-Ig at 15mg/kg per mouse an effect was observed [43].

We demonstrated that the superiority of RBD-Ig was not due to quantitative or qualitative changes in the antibody response, we thus hypothesized that it may be due to RBD-Ig effect on its target protein ACE2. To check this, we first wanted to assess whether changes occur in ACE2 surface expression as it is targeted by RBD-Ig. No changes were observed in ACE2 surface expression in SARS-CoV-2 infected cells at different time points. Although many suggest that an ACE2 downregulation might occur during infection [44][45][46], to the best of our knowledge this has not been investigated, perhaps because there was no effective commercial antibody available against ACE2. We also assessed ACE2 surface expression following RBD-Ig incubation and saw that ACE2 expression did not change drastically. Another important check was of ACE2 enzymatic activity following binding to SARS-CoV-2 RBD as it was reported to enhance ACE2 activity [47]. In contrast, we report here that ACE2 activity was not affected following incubation with RBD-Ig. The reason for this discrepancy is not understood. We next hypothesized that RBD-Ig blocks infection by physically interacting with ACE2. We further thought that RBD-Ig is more efficient than ACE2-Ig since RBD-Ig, binds to the constantly expressed ACE2 on the target cells, while ACE2-Ig interacts with the spike protein found on a replicating virus. Indeed, we showed that RBD-Ig can neutralizes in vitro SARS-CoV-2 even at high virus titers, while ACE2-Ig cannot.

To summarize we suggest that RBD-Ig inhibit SARS-CoV-2 infection by physically blocking ACE2. Thus, RBD-Ig is particularly advantageous as a treatment for SARS-CoV-2 infection since it targets ACE2 which expression on cell surface remains almost constant, rather than a mutating and replicating virus.

## Methods

### Cell lines and viruses

293T cells (CRL-3216) were grown in Dulbecco's modified Eagle's medium (DMEM, Sigma-Aldrich) containing 10% Fetal bovine serum (FBS), (Sigma-Aldrich), 1% L-glutamine (Biological Industries (BI)), 1% sodium pyruvate (BI), 1% nonessential amino acids (BI), and 1% penicillin-streptomycin (BI). Vero E6 cells (CRL-1586) were grown in DMEM containing 10% FBS, MEM non-essential amino acids (NEAA), 2mM L-Glutamine, 100Units/ml Penicillin, 0.1mg/ml streptomycin, 12.5 Units/ml Nystatin (P/S/N) (BI). All cells were cultured at 37°C, 5% CO2 at 95% air atmosphere. SARS-CoV-2 (GISAID accession EPI_ISL_406862) was kindly provided by Bundeswehr Institute of Microbiology, Munich, Germany. Virus stocks were propagated (4 passages) and tittered on Vero E6 cells. Handling and experiments with SARS-CoV-2 virus were conducted in a BSL3 facility in accordance with the biosafety guidelines of the Israel Institute for Biological Research (IIBR).

### Mice

Homozygous female outbred K18-hACE2 transgenic mice (2B6.Cg-Tg(K18-ACE2)2Prlmn/J, Stock No: 034860, Jackson laboratory) 6-8 weeks old were maintained at 20-22°C with relative humidity of 50 ± 10% on a 12hrs light/dark cycle. Animals were fed with commercial rodent chow (Koffolk Inc.) and provided with tap water ad libitum. Prior infection, mice were kept in groups of 10. Mice were randomly assigned to experimental groups of 7-8 mice per group. 200 PFU of SARS-CoV-2 (10-15 LD50) was diluted in PBS supplemented with 2% FBS (BI) to infect animal by 20µl intranasal instillation of anesthetized mice. Body weight was monitored daily over 13-15 days. At 15 dpi mice were bled through the venous tail and sera were obtained. Residual SARS-CoV-2 virus in the sera was neutralized by heating to 60°C for 30 minutes. Four groups of mice were used: 1. Naïve (uninfected & untreated mice). 2. Infected and treated with Control-Ig. 3. Infected and treated with ACE2-Ig. 4. Infected and treated with RBD-Ig.

### Flow cytometry

Primary antibody staining was performed at 4°C for 1 hour, cells were then washed in FACS buffer (1% BSA and 0.05% Sodium Azide in phosphate-buffered saline) and secondary antibody was added for 30 minutes at 4°C. Then, cells were washed in FACS buffer and fixed with 4% paraformaldehyde for 20 minutes followed by CytoFlex analysis. We used the following primary antibodies: Rabbit MAb SARS-CoV-2 Spike S1 Antibody (Cat#40150-R007-100, Sino Biological), Purified anti-DYKDDDDK Tag Antibody (Cat#637302, BioLegend), anti-ACE2 01 (generated by us). The following secondary antibodies were used: Alexa Fluor 647-conjugated Goat Anti-Rabbit IgG (Cat#111-606-144, Jackson ImmunoResearch Laboratories), Alexa Fluor 647-conjugated Donkey anti-human IgG (Cat#709-606-098, Jackson ImmunoResearch Laboratories), Alexa Fluor 647-conjugated Goat Anti-Mouse IgG (Cat#115-606-062, Jackson ImmunoResearch Laboratories). Data were analyzed using FCS Express 6/7.

### Fusion proteins

PCR-generated fragments encoding the extracellular part of human ACE2 or SARS-CoV-2 RBD were each cloned into vectors containing the Fc portion of human IgG1, and a Puromycin resistance gene. Sequencing of the constructs revealed that cDNA of all Ig-fusion proteins was in frame with the human Fc genomic DNA and were identical to the reported sequences. The Ig-vectors were then introduced to 293T cells (CRL-3216, ATCC) and the transfected cells were grown in the continuous presence of Puromycin. The ACE2-Ig and RBD-Ig fusion proteins secreted to the medium were purified on HiTrap Protein G High Performance column (Cat#GE17-0405-01, GE Healthcare). Control-Ig was one of the following fusion proteins: KIR2DL1-Ig/KIR2DS1-Ig/ CD59-Ig/CD16-Ig, which were previously made in our lab as described here [48]. RBD PCR-generated fragments were made from 2 separated PCR reactions followed by a third reaction in which we used the forward primer of reaction 1, the reverse primer of reaction 2 and the products from reaction 1 and 2 as a template. The RBD portion of the fusion protein is composed of 331-524 AA from the full spike protein fused to the IgG1 human portion. Primer FW for ACE2-Ig:
AAAGCTAGCGCCGCCACCATGTCAAGCTCTTCCTGGC. Primer RV for ACE2-Ig:
TTTTGATCAGAAACAGGGGGCTG. Primer FW for RBD-Ig reaction 1:
AAATTGAATTCGCCGCCACCATGCCCATGGGGTCTCTGCA. Primer RV for RBD-Ig reaction 1: GTTGGTGATGTTTCCGAGGCAGGAAGCGACC. Primer FW for RBD-Ig reaction 2: GCCTCGGAAACATCACCAACCTGTGTCCAT. Primer RV for RBD-Ig reaction 2: TTTGGATCCACTGTGGCAGGGGCATGG.

### Lentivirus production

Lentiviral vectors were produced by transient three-plasmid transfection as described here [49]. First, 293T cells were grown overnight in 6-well plates (2.2X10^5^ cells/well). The following day pMD.G / VSV-G/ SARS-CoV-2 spike envelope expressing plasmid (0.35 μg/well), a gag-pol packaging construct (0.65 μg/well) and the relevant vector construct (1 μg/well) were transfected using the TransIT®-LT1 Transfection Reagent (MIR 2306, Mirus). Two days after transfection the soups containing the viruses were collected and filtered.

### Generation of 293T-ACE2 cells

ACE2 was amplified from cDNA and an N-terminal Flag-Tag was introduced immediately after the signal peptide. The flag-tagged ACE2 was cloned into the plasmid pHAGE-DsRED(-) GFP(+). This plasmid carrying the Flag-tagged ACE2 was used as a vector construct to produce lentiviruses as described above. The resulting lentiviruses were used to infect 293T cells. The transduced cells were stained with anti-human ACE-2, RBD-Ig and checked for GFP percentage by Flow Cytometry. PCR-generated fragments were made from 2 separated PCR reactions followed by a third reaction in which we used the forward primer of reaction 1, the reverse primer of reaction 2 and the products from reaction 1 and 2 as a template. Primer FW reaction 1:
AAATTGAATTCGCCGCCACCATGCCCATGGGGTCTCTGCA. Primer RV reaction 1: GTTGGTGATGTTTCCGAGGCAGGAAGCGACC. Primer FW reaction 2: GCCTCGGAAACATCACCAACCTGTGTCCAT. Primer RV reaction 2: TTTGGATCCACTGTGGCAGGGGCATGG.

### Generation of 293T-Spike cells

First, 293T cells were grown overnight in 6-well plates (2.2X10^5^ cells/well). Then SARS-CoV-2 spike envelope expression plasmid was co-transfected as described above with the plasmid pHAGE-DsRED(-) GFP(+) as a vector construct. As a control we performed the same co-transfection but with the VSV-G envelope plasmid. 48 hours following transfection, media (containing lentiviruses) was removed, and cells were used for flow cytometry experiments. Transfection efficiency was assessed by GFP expression. For each flow cytometry experiment we generated new 293T-Spike cells as described here.

### Enzymatic activity

The enzymatic activity of the ACE2-Ig fusion protein was evaluated using the ACE2 Activity Assay Kit (Fluorometric) (Cat#BN01071, Assay Genie) according to the manufacturer instructions. 0.8 ug/well of ACE2-Ig was used with or without the inhibitor supplied with the kit. The 293T-ACE2 cells lysate was prepared and 10 ug of it was incubated with RBD-Ig according to the manufacturer instructions. Plates were read by Tecan Spark 10M and data were analyzed using Magellan 1.1.

### Fusion protein staining with inhibitor

0.8 ug/well of ACE2-Ig was incubated with or without the ACE2 inhibitor (supplied with the kit described above) for 15 minutes at room temperature. Then, ACE2-Ig (with or without the inhibitor) was added to either the 293T parental cells or to the 293T-Spike cells for 1 hour at 4°C. Afterwards, cells were washed in FACS buffer and stained with Alexa Fluor 647-conjugated anti-human IgG secondary antibody. Then, cells were washed in FACS buffer and analyzed by CytoFlex.

### SARS-CoV-2 Plaque reduction neutralization test (PRNT) with ACE2-Ig

Vero E6 cells (CRL-1586, ATCC) were seeded in 12-well plates (5×10^5^ cells/well) and grown overnight in Penicillin-Streptomycin-Neomycin (P/S/N, BI) containing medium. The following day, ACE2-Ig and Control-Ig were either diluted to 50µg/ml-0.048µg/ml or 200ug/ml in 400µl of MEM containing 2% FBS, NEAA, 2mM L-Glutamine, and P/S/N. The diluted fusion proteins ACE2-Ig and Control-Ig were then mixed with 400µl of 300 PFU (Plaque Forming Units)/ml or 100-218,700 PFU/ml of SARS-CoV-2. The virus-protein mixtures were incubated at 37°C, 5% CO2 for 1 hour. Vero E6 cell monolayers were washed once with DMEM and 200µl of each dilution of protein-virus mixture was added in triplicates for 1 hour at 37°C. Virus without fusion protein served as control. 2ml/well overlay {MEM containing 2% FBS and 0.4% Tragacanth (Sigma-Aldrich)} were added to each well and plates were incubated at 37°C 5% CO2 for 48 hours. The overlay was then aspirated, the cells were fixed and stained with 1ml of crystal violet solution (BI). The number of plaques in each well were determined and neutralization percentages were calculated as follows: 100 × [1 – (average number of plaques for each dilution/average number of the virus dose control plaques)]. SARS-CoV-2 strain used was kindly provided by Bundeswehr Institute of Microbiology, Munich, Germany (GISAID accession EPI_ISL_406862)

### SARS-CoV-2 PRNT with RBD-Ig

Vero E6 cells were seeded in 12-well plates as described above. The next day, RBD-Ig and Control-Ig were either diluted to 25µg/ml-0.024µg/ml or 100ug/ml in 400µl of MEM containing 2% FBS, NEAA, 2mM L-Glutamine, and P/S/N. Cell monolayers were washed once with DMEM and the diluted fusion protein RBD-Ig or Control-Ig was then added in triplicates (200µl/well). Cell monolayers were then incubated at 37°C, 5% CO2 for 1 hour. Afterwards 100µl of 300 PFU (Plaque Forming Units)/ml or 100-218,700 PFU/ml of SARS-CoV-2 was added for 1 hour at 37°C. Then 2ml/well overlay were added, and plates were incubated at 37°C 5% CO2 for 48 hours, as described above. The cells were then fixed and stained, and neutralization percentages were determined as described above.

### In vivo treatment with fusion proteins

SARS-CoV-2 infected mice were treated with 75ug/mouse of the fusion protein (Control-Ig/ ACE2-Ig/ RBD-Ig) at 3 time points: day 1, day 2/3 and day 3/5 post-infection (PI). Treatment was intraperitoneally (IP) administered in 300 ul. Mice were infected with a SARS-CoV-2 strain kindly provided by Prof. Dr. Christian Drosten (Charité, Berlin) (EVAg Ref-SKU: 026V-03883).

### Staining with mice sera

Sera were obtained 15 dpi from the various immunized groups and from naïve mice. Sera were diluted to 1:500, 1:1K, 1:5K, 1:10K per well and added to 50,000 293T-Parental cells or 293T-Spike cells in a 96-U-well plate for 1 hour at 4°C. Cells were then washed, and an Alexa Fluor 647 Anti-Mouse IgG secondary antibody was added.

### Blocking with mice sera

Sera from the various immunized mice groups was diluted to 1:100 per well and added to 50K 293T-Parental cells or 293T-Spike cells in a 96-U-well plate for 1 hour at 4°C. Afterwards, ACE2-Ig was added as a primary antibody for 1 hour at 4°C. Then, cells were washed, and Alexa Fluor 647 Anti-human IgG secondary antibody was added.

### Statistics

Statistical analysis were performed using either Prism 8 (GraphPad) or Excel (Microsoft). Error bars represent SD. All the relevant statistical data for the experiments including the statistical test used, value of n, definition of significance, etc. can be found in the figure legends or the relevant method section.

### Study approval

Animal experiments involving SARS-CoV-2 were conducted in a BSL3 facility and treatment of animals was in accordance with regulations outlined in the U.S. Department of Agriculture (USDA) Animal Welfare Act and the conditions specified in the Guide for Care and Use of Laboratory Animals (National Institute of Health, 2011). Animal studies were approved by the local IIBR ethical committee on animal experiments (protocol number M-54-20).

## Author contributions

Conceptualization, O.M. and A.C.; Methodology, O.M, A.C., H.A., I.K., D.W.; Investigation, A.C., H.A., I.K., I.B., T.L.R., G.R., O.A, E.B.V., T.I., S.M., B.P., E.Z.; Resources, O.M., H.A., E.B.V., T.I., S.M., B.P., E.Z., D.W., S.J., O.A.; Writing – Original Draft, O.M. and A.C.; Writing – Review & Editing, O.M., A.C., S.J. and O.B.; Visualization, O.B.; Supervision, O.M.; Project Administration, O.M.; Funding Acquisition, O.M., I.B. and S.J.

## Acknowledgments

The authors would like to thank Prof. Dr. Christian Drosten at the Charité-Universitätsmedizin, Institute of Virology, Berlin, Germany for providing the SARS-CoV-2 BavPat1/2020 strain. We would like to also thank Dr. Alex Rouvinski (Hebrew university, Israel) for kindly providing the SARS-CoV-2 spike envelope plasmid. Figure 1A was generated using BioRender.com. This work was supported by Integra Holdings, the Israel Science Foundation (Moked grant), the GIF Foundation, the ICRF professorship grant, the ISF Israel- China grant, the MOST-DKFZ grant, the ERC Marie Currie grant, the Rothschild Foundation, the Croatian Science Foundation (IP-CORONA-04-2073) (I.B.) and by the grant “Strengthening the capacity of CerVirVac for research in virus immunology and vaccinology”, KK.01.1.1.01.0006, awarded to the Scientific Centre of Excellence for Virus Immunology and Vaccines and co-financed by the European Regional Development Fund (S.J.).

## References

1. Chakraborty I, Maity P. COVID-19 outbreak: Migration, effects on society, global environment and prevention. Sci Total Environ [Internet]. 2020 Aug 1 [cited 2020 Nov 22];728. Available from: https://doi.org/10.1016/j.scitotenv.2020.138882

2. Administration D. Pfizer COVID-19 Vaccine EUA Letter of Authorization reissued 12-23-20. 2020.

3. Administration D. Moderna COVID-19 Vaccine EUA Letter of Authorization. 2020.

4. A Study of Ad26.COV2.S for the Prevention of SARS-CoV-2-Mediated COVID-19 in Adult Participants - Full Text View - ClinicalTrials.gov [Internet]. [cited 2021 Apr 6]. Available from: https://clinicaltrials.gov/ct2/show/NCT04505722#wrapper

5. Polack FP, Thomas SJ, Kitchin N, Absalon J, Gurtman A, Lockhart S, et al. Safety and Efficacy of the BNT162b2 mRNA Covid-19 Vaccine. N Engl J Med [Internet]. 2020 Dec 31 [cited 2021 Feb 8];383(27):2603–15. Available from: http://www.nejm.org/doi/10.1056/NEJMoa2034577

6. Baden LR, El Sahly HM, Essink B, Kotloff K, Frey S, Novak R, et al. Efficacy and Safety of the mRNA-1273 SARS-CoV-2 Vaccine. N Engl J Med [Internet]. 2021 Feb 4 [cited 2021 Feb 8];384(5):403–16. Available from: http://www.nejm.org/doi/10.1056/NEJMoa2035389

7. Williams TC, Burgers WA. SARS-CoV-2 evolution and vaccines: cause for concern? Lancet Respir Med [Internet]. 2021 Jan 29 [cited 2021 Feb 4];0(0). Available from: http://www.ncbi.nlm.nih.gov/pubmed/33524316

8. Wang Q, Zhang Y, Wu L, Niu S, Song C, Zhang Z, et al. Structural and Functional Basis of SARS-CoV-2 Entry by Using Human ACE2. Cell [Internet]. 2020 May 14 [cited 2020 Aug 11];181(4):894–904.e9. Available from: /pmc/articles/PMC7144619/?report=abstract

9. Shang J, Ye G, Shi K, Wan Y, Luo C, Aihara H, et al. Structural basis of receptor recognition by SARS-CoV-2. Nature [Internet]. 2020 May 14 [cited 2020 Nov 18];581(7807):221–4. Available from: /pmc/articles/PMC7328981/?report=abstract

10. loganathan S, Kuppusamy M, Wankhar W, Gurugubelli KR, Mahadevappa VH, Lepcha L, et al. Angiotensin-converting enzyme 2 (ACE2): COVID 19 gate way to multiple organ failure syndromes. Vol. 283, Respiratory Physiology and Neurobiology. Elsevier B.V.; 2021. p. 103548.

11. Wang T, Du Z, Zhu F, Cao Z, An Y, Gao Y, et al. Comorbidities and multi-organ injuries in the treatment of COVID-19. Lancet [Internet]. 2020 [cited 2020 Nov 25];395:e52. Available from: https://www.bbc.co.

12. Wang W, McKinnie SMK, Farhan M, Paul M, McDonald T, McLean B, et al. Angiotensin-Converting Enzyme 2 Metabolizes and Partially Inactivates Pyr-Apelin-13 and Apelin-17: Physiological Effects in the Cardiovascular System. Hypertension [Internet]. 2016 Aug 1 [cited 2020 Nov 18];68(2):365–77. Available from: https://www.ahajournals.org/doi/10.1161/HYPERTENSIONAHA.115.06892

13. Flores-Muñoz M, Smith NJ, Haggerty C, Milligan G, Nicklin SA. Angiotensin1-9 antagonises pro-hypertrophic signalling in cardiomyocytes via the angiotensin type 2 receptor. J Physiol [Internet]. 2011 Feb [cited 2020 Nov 18];589(4):939–51. Available from: https://pubmed.ncbi.nlm.nih.gov/21173078/

14. Donoghue M, Hsieh F, Baronas E, Godbout K, Gosselin M, Stagliano N, et al. A novel angiotensin-converting enzyme-related carboxypeptidase (ACE2) converts angiotensin I to angiotensin 1-9. Circ Res [Internet]. 2000 Sep 1 [cited 2020 Nov 18];87(5). Available from: http://www.circresaha.org.

15. Rebold N, Holger D, Alosaimy S, Morrisette T, Rybak M. COVID-19: Before the Fall, An Evidence-Based Narrative Review of Treatment Options. Infect Dis Ther [Internet]. 2021 Jan 25 [cited 2021 Feb 4];1–21. Available from: http://www.ncbi.nlm.nih.gov/pubmed/33495967

16. Beck A, Reichert JM. Therapeutic Fc-fusion proteins and peptides as successful alternatives to antibodies [Internet]. Vol. 3, mAbs. MAbs; 2011 [cited 2020 Nov 18]. p. 415–6. Available from: https://pubmed.ncbi.nlm.nih.gov/21785279/

17. Capon DJ, Chamow SM, Mordenti J, Marsters SA, Gregory T, Mitsuya H, et al. Designing CD4 immunoadhesins for AIDS therapy. Nature. 1989;337(6207):525–31.

18. Czajkowsky DM, Hu J, Shao Z, Pleass RJ. Fc-fusion proteins: New developments and future perspectives [Internet]. Vol. 4, EMBO Molecular Medicine. EMBO Mol Med; 2012 [cited 2020 Nov 18]. p. 1015–28. Available from: https://pubmed.ncbi.nlm.nih.gov/22837174/

19. Lei C, Qian K, Li T, Zhang S, Fu W, Ding M, et al. Neutralization of SARS-CoV-2 spike pseudotyped virus by recombinant ACE2-Ig. Nat Commun [Internet]. 2020 Dec 1 [cited 2020 Jun 30];11(1):1–5. Available from: https://www.nature.com/articles/s41467-020-16048-4

20. Monteil V, Kwon H, Prado P, Hagelkrüys A, Wimmer RA, Stahl M, et al. Inhibition of SARS-CoV-2 Infections in Engineered Human Tissues Using Clinical-Grade Soluble Human ACE2. Cell. 2020 May 14;181(4):905–913.e7.

21. Liu X, Drelich A, Li W, Chen C, Sun Z, Shi M, et al. Enhanced elicitation of potent neutralizing antibodies by the SARS-CoV-2 spike receptor binding domain Fc fusion protein in mice. Vaccine. 2020 Oct 27;38(46):7205–12.

22. Quinlan BD, Mou H, Zhang L, Guo Y, He W, Ojha A, et al. The SARS-CoV-2 Receptor-Binding Domain Elicits a Potent Neutralizing Response Without Antibody-Dependent Enhancement. SSRN Electron J [Internet]. 2020 Apr 15 [cited 2020 Nov 18]; Available from: https://papers.ssrn.com/abstract=3575134

23. Xiao L, Sakagami H, Miwa N. ACE2: The key molecule for understanding the pathophysiology of severe and critical conditions of COVID-19: Demon or angel? [Internet]. Vol. 12, Viruses. MDPI AG; 2020 [cited 2020 Nov 18]. Available from: https://pubmed.ncbi.nlm.nih.gov/32354022/

24. Devaux CA, Rolain JM, Raoult D. ACE2 receptor polymorphism: Susceptibility to SARS-CoV-2, hypertension, multi-organ failure, and COVID-19 disease outcome. Vol. 53, Journal of Microbiology, Immunology and Infection. Elsevier Ltd; 2020. p. 425–35.

25. Zhou P, Yang X Lou, Wang XG, Hu B, Zhang L, Zhang W, et al. A pneumonia outbreak associated with a new coronavirus of probable bat origin. Nature [Internet]. 2020 Mar 12 [cited 2021 Feb 8];579(7798):270–3. Available from: https://doi.org/10.1038/s41586-020-2012-7

26. Winkler ES, Bailey AL, Kafai NM, Nair S, McCune BT, Yu J, et al. SARS-CoV-2 infection of human ACE2-transgenic mice causes severe lung inflammation and impaired function. Nat Immunol [Internet]. 2020;21(11):1327–35. Available from: http://dx.doi.org/10.1038/s41590-020-0778-2

27. Ju B, Zhang Q, Ge J, Wang R, Sun J, Ge X, et al. Human neutralizing antibodies elicited by SARS-CoV-2 infection. Nature [Internet]. 2020 Aug 6 [cited 2020 Nov 25];584(7819):115–9. Available from: https://pubmed.ncbi.nlm.nih.gov/32454513/

28. World Health Organization. COVID-19 Weekly Epidemiological Update 22. World Heal Organ [Internet]. 2021;(January):1–3. Available from: https://www.who.int/docs/default-source/coronaviruse/situation-reports/weekly_epidemiological_update_22.pdf

29. Cohen J. Vaccine wagers on coronavirus surface protein pay off. Science (80-) [Internet]. 2020 Nov 20 [cited 2020 Nov 30];370(6519):894–5. Available from: https://www.sciencemag.org/lookup/doi/10.1126/science.370.6519.894

30. Tegally H, Wilkinson E, Giovanetti M, Iranzadeh A, Fonseca V, Giandhari J, et al. Sibongile Walaza 9. Arghavan Alisoltani-Dehkordi [Internet]. 2020 Dec 22 [cited 2021 Feb 5];10:2020.12.21.20248640. Available from: https://doi.org/10.1101/2020.12.21.20248640

31. Liu Z, Zheng H, Lin H, Li M, Yuan R, Peng J, et al. Identification of Common Deletions in the Spike Protein of Severe Acute Respiratory Syndrome Coronavirus 2. J Virol [Internet]. 2020 Jun 22 [cited 2021 Feb 5];94(17):790–810. Available from: http://jvi.asm.org/

32. Plante JA, Liu Y, Liu J, Xia H, Johnson BA, Lokugamage KG, et al. Spike mutation D614G alters SARS-CoV-2 fitness. Nature [Internet]. 2020 Oct 26 [cited 2021 Feb 5];1–6. Available from: https://doi.org/10.1038/s41586-020-2895-3

33. Volz E, Mishra S, Chand M, Barrett JC, Johnson R, Hopkins S, et al. Transmission of SARS-CoV-2 Lineage B.1.1.7 in England: Insights from linking epidemiological and genetic data. medRxiv [Internet]. 2021 Jan 4 [cited 2021 Feb 5];2020.12.30.20249034. Available from: https://www.medrxiv.org/content/10.1101/2020.12.30.20249034v2

34. Jangra S, Ye C, Rathnasinghe R, Stadlbauer D, Krammer F, Simon V, et al. The E484K mutation in the SARS-CoV-2 spike protein reduces but does not abolish neutralizing activity of human convalescent and post-vaccination sera. medRxiv Prepr Serv Heal Sci [Internet]. 2021 Jan 29 [cited 2021 Feb 4];2021.01.26.21250543. Available from: http://www.ncbi.nlm.nih.gov/pubmed/33532796

35. Wibmer CK, Ayres F, Hermanus T, Madzivhandila M, Kgagudi P, Lambson BE, et al. SARS-CoV-2 501Y.V2 escapes neutralization by South African COVID-19 donor plasma. bioRxiv [Internet]. 2021 Jan 19 [cited 2021 Feb 5];2021.01.18.427166. Available from: https://doi.org/10.1101/2021.01.18.427166

36. Gottlieb RL, Nirula A, Chen P, Boscia J, Heller B, Morris J, et al. Effect of Bamlanivimab as Monotherapy or in Combination With Etesevimab on Viral Load in Patients With Mild to Moderate COVID-19. JAMA [Internet]. 2021 Jan 21 [cited 2021 Feb 5]; Available from: https://jamanetwork.com/journals/jama/fullarticle/2775647

37. Yang P, Gu H, Zhao Z, Wang W, Cao B, Lai C, et al. Angiotensin-converting enzyme 2 (ACE2) mediates influenza H7N9 virus-induced acute lung injury. Sci Rep [Internet]. 2014 Nov 13 [cited 2020 Nov 18];4(1):7027. Available from: https://www.nature.com/scientificreports

38. Zou Z, Yan Y, Shu Y, Gao R, Sun Y, Li X, et al. Angiotensin-converting enzyme 2 protects from lethal avian influenza A H5N1 infections. Nat Commun [Internet]. 2014 May 6 [cited 2020 Nov 18];5(1):1–7. Available from: https://www.nature.com/naturecommunications

39. Wang X, Khaidakov M, Ding Z, Mitra S, Lu J, Liu S, et al. Cross-talk between inflammation and angiotensin II: Studies based on direct transfection of cardiomyocytes with AT1R and AT2R cDNA. Exp Biol Med [Internet]. 2012 Dec [cited 2020 Nov 18];237(12):1394–401. Available from: https://www.pubmed.ncbi.nlm.nih.gov/23354398/

40. Sodhi CP, Wohlford-Lenane C, Yamaguchi Y, Prindle T, Fulton WB, Wang S, et al. Attenuation of pulmonary ACE2 activity impairs inactivation of des-arg9 bradykinin/BKB1R axis and facilitates LPS-induced neutrophil infiltration. Am J Physiol - Lung Cell Mol Physiol [Internet]. 2018 Jan 1 [cited 2020 Nov 18];314(1):L17–31. Available from: https://pubmed.ncbi.nlm.nih.gov/28935640/

41. Moore MJ, Dorfman T, Li W, Wong SK, Li Y, Kuhn JH, et al. Retroviruses Pseudotyped with the Severe Acute Respiratory Syndrome Coronavirus Spike Protein Efficiently Infect Cells Expressing Angiotensin-Converting Enzyme 2. J Virol [Internet]. 2004 Oct 1 [cited 2020 Aug 11];78(19):10628–35. Available from: http://jvi.asm.org/

42. Huang K, Lin M, Kuo T, Chen C, Lin C, Chou Y, et al. Humanized COVID‐ 19 decoy antibody effectively blocks viral entry and prevents SARS‐ CoV‐ 2 infection. EMBO Mol Med [Internet]. 2021 Jan 11 [cited 2021 Apr 4];13(1). Available from: https://pubmed.ncbi.nlm.nih.gov/33159417/

43. Iwanaga N, Cooper L, Rong L, Beddingfield B, Crabtree J, Tripp RA, et al. Novel ACE2-IgG1 fusions with improved in vitro and in vivo activity against SARS-CoV2. [cited 2021 Apr 6]; Available from: https://doi.org/10.1101/2020.06.15.152157

44. Patel SK, Juno JA, Lee WS, Wragg KM, Hogarth PM, Kent SJ, et al. Plasma ACE2 activity is persistently elevated following SARS-CoV-2 infection: implications for COVID-19 pathogenesis and consequences. Eur Respir J [Internet]. 2021 Jan 21 [cited 2021 Feb 5];2003730. Available from: http://erj.ersjournals.com/lookup/doi/10.1183/13993003.03730-2020

45. Ciulla MM. SARS-CoV-2 downregulation of ACE2 and pleiotropic effects of ACEIs/ARBs [Internet]. Vol. 43, Hypertension Research. Springer Nature; 2020 [cited 2021 Feb 5]. p. 985–6. Available from: https://doi.org/10.1038/s41440-020-0488-z

46. Zhang P, Zhu L, Cai J, Lei F, Qin JJ, Xie J, et al. Association of Inpatient Use of Angiotensin-Converting Enzyme Inhibitors and Angiotensin II Receptor Blockers with Mortality among Patients with Hypertension Hospitalized with COVID-19. Circ Res [Internet]. 2020 Jun 5 [cited 2021 Feb 5];126(12):1671–81. Available from: https://www.ahajournals.org/doi/suppl/10.1161/CIRCRESAHA.120.317134.

47. Lu J, Sun P. High affinity binding of SARS-CoV-2 spike protein enhances ACE2 carboxypeptidase activity. bioRxiv Prepr Serv Biol [Internet]. 2020 [cited 2021 Feb 5]; Available from: /pmc/articles/PMC7337377/?report=abstract

48. Mandelboim O, Malik P, Davis DM, Jo CH, Boyson JE, Strominger JL. Human CD16 as a lysis receptor mediating direct natural killer cell cytotoxicity. Proc Natl Acad Sci U S A [Internet]. 1999 May 11 [cited 2020 Nov 19];96(10):5640–4. Available from: https://www.pnas.org.

49. Naldini L, Blömer U, Gallay P, Ory D, Mulligan R, Gage FH, et al. In vivo gene delivery and stable transduction of nondividing cells by a lentiviral vector. Science (80-) [Internet]. 1996 Apr 12 [cited 2020 Nov 19];272(5259):263–7. Available from: https://www.pubmed.ncbi.nlm.nih.gov/8602510/

